# Dissecting the role of RNA-binding proteins in early herpes simplex virus 1 transcription using acute protein depletion

**DOI:** 10.1101/2025.02.10.637450

**Authors:** Niclas Barke, Sawinee Masser, Doerte Stalling, Emanuel Wyler, Robert Patrick Zinzen, Sabrina Schreiner, Jens Bosse, Markus Landthaler

## Abstract

Herpes simplex virus 1 (HSV-1) infects about 50-80% of the entire human population and persists in the neurons of affected individuals. A fraction of affected individuals suffer from recurrent cold sores caused by reactivating virus, in rare but severe cases the virus can cause encephalitis.

During lytic infection, the virus relies on host factors such as RNA polymerase II and accessory proteins involved in transcription to express its genes and ensure successful replication. In general, RNA molecules in cells are bound by RNA-binding proteins (RBPs) during their entire lifecycle. Importantly, RBPs are increasingly described to also regulate transcription, an aspect long time outside the scope of investigations, especially during viral infections. Here, we examined the impact of five nuclear proteins (FUBP1, SLBP, SFPQ, SPT5 and SAF-B) with known RNA-binding activities on HSV-1 transcription. Additionally, we evaluated their importance for human adenovirus C5 (HAdV) growth to assess whether these host factors are specific to HSV-1 infections or might have broader relevance for the general transcription of dsDNA viruses.

We show that the transcriptional elongation factor SPT5 coded by SUPT5H accumulates on HSV-1 genomes early during the infection and is required for the transcription of the immediate-early gene UL54. Its depletion affects also HAdV replication, indicating a general role in transcription of viruses that depend on the host transcriptional machinery.

In contrast, depletion of the transcriptional repressor and paraspeckle protein SFPQ reduces UL54 RNA levels in HSV-1 infection, but does not cause significant changes in HAdV growth. Since SFPQ does not co-localize with HSV-1 genomes, this suggests a function not directly associated to viral DNA.

## Introduction

Herpes simplex virus 1 (HSV-1) is transmitted between humans by direct contact and establishes lifelong latency in neurons of the trigeminal ganglia^1,2^. As of 2012, an estimated 3.7 billion people under the age of 49 were infected with HSV-1 worldwide^3^. While some individuals experience recurrent painful blisters and sores, typically at the lips following viral reactivation, HSV-1 can lead to severe complications such as keratitis, blindness and encephalitis^1,4,5^. Newborns, the elderly and immunocompromised individuals are particularly vulnerable to these outcomes^5^. Furthermore, growing evidence suggests a contribution of HSV-1 infections and/or reactivations to the development and/or progression of neurodegenerative diseases, including Alzheimer’s^6–8^. Current treatment options mainly rely on inhibitors and nucleoside analogs such as acyclovir, which are susceptible to viral resistance highlighting the urgent need for alternative drugs^5,9^. Despite ongoing research, an effective vaccine remains unaivalable^9^. As a result, HSV-1 continues to pose a global health concern. Understanding and targeting the early stages of the infection may offer new opportunities to prevent and ultimately reduce this burden.

To initiate HSV-1 transcription mediated by the RNA polymerase II (Pol II), the tegument protein VP16 functions as an activator working together with the host transcription factors (TFs) HCF and Oct-1 to drive expression of the 5 immediate-early genes: RL2 (ICP0), RS1 (ICP4), US1 (ICP22), UL54 (ICP27) and US12 (ICP47)^10–13^. Among these, ICP4 plays a crucial role in orchestrating the viral expression cascade by interacting with the general host transcription factors of mediator and components of the TFIID complex, to activate transcription of early and late genes^14–19^. It exemplifies that despite its relatively large genome of about 152 kbp, HSV-1 relies on the activity of several host factors to facilitate viral transcription and replication^20^. This includes host RNA-binding proteins (RBPs), that associate with viral RNA (vRNA) throughout the entire life cycle from nuclear transcription to translation in the cytoplasm^21,22^. While classical functions assigned to RBPs include mRNA splicing, modification, stability, localization, and translation, in recent years emerging evidence suggests that some RBPs also contribute to transcriptional regulation, a function that remains poorly understood in the context of viral infections^23,24^. While RBPs are crucial for various aspects of RNA metabolism, most virological studies have primarily focused on their involvement in innate immune recognition and decay of vRNAs, as well as their role during the intracellular transport, nuclear import/export and translation of vRNAs. Many RBPs are indispensable for cellular functions, making knockout approaches challenging. Since specific inhibitors are generally unavailable, depletion strategies often rely on CRISPR/Cas9 or RNA interference (RNAi). However, RNAi leads to a gradual decrease of protein levels over one to three days, increasing the risk of observing secondary effects such as reduced cell viability.

To address this issue and elucidate the role of RBPs in HSV-1 infection, we used an auxin-mediated protein degradation system^25^. This approach enables rapid depletion of target proteins within 2-6 hours by harnessing cellular E3-ubiquitin ligases and subsequent proteasomal degradation. Unlike RNAi, this method allows us to assess the direct, immediate impact of these proteins on the viral replication cycle, as the short depletion time reduces the possibility of secondary effects. We endogenously degron-tagged several RBPs involved in transcriptional regulation and previously identified by an RNAi screen to influence vRNA levels at early times of infection and vDNA replication^25^. Our findings indicate that the human transcription elongation factor SUPT5H, is associated with the vDNA at early times of infection and is likely essential for the transcription of HSV-1 genes. Furthermore, we demonstrate that the absence of SFPQ, a multifunctional protein involved in splicing and paraspeckle formation, leads to a reduction of vRNA levels but does not completely block transcription^26–29^. To facilitate comparative analysis, we also included adenovirus C5, another double-stranded DNA virus replicating in the nucleus in our study. Despite its genome being only a quarter to a third the size of HSV-1, they share a broadly similar replication strategy^30^. This comparative approach allows us to distinguish factors specific to HSV-1 from those with more general functions for dsDNA viruses replicating in the nucleus.

## Material and Methods

### Cell lines and culturing conditions

HCT116 (ATCC, CCL-247) cells were cultured in McCoy’s 5A (Modified) Medium (ThermoFisher) with 2 mM L-glutamine (ThermoFisher) and 9-10% fetal bovine serum (ThermoFisher). HaCaT cells for EdC-labeled virus stock production were kept in DMEM (high glucose) (Sigma-Aldrich) supplemented with GlutaMAX and 10% FBS. N- and C-termini of genes of interest were tagged with miniIAA7(AID) in F-box protein AtAFB2 expressing HCT116 cells as described in Boehm et al.^31^. Used sgRNAs are listed in Table 1. Cells were maintained at 37°C and 5% CO2 in a humidified incubator.

**Table 1:**
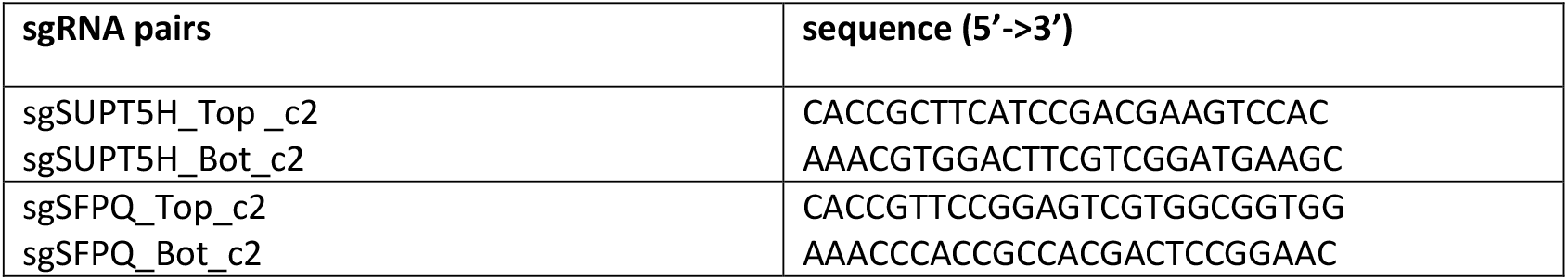
List of sgRNAs used to AID-tag HCT116 cells by CRISPR/Cas9 genome editing.

A549 cells (ATCC, CCL-185) and HEK293 (ATCC, CRL-1573) were grown in Dulbecco’s modified Eagle’s medium (DMEM) supplemented with 5% fetal calf serum, 100 U of penicillin, 100 μg of streptomycin per ml in a 5% CO2 atmosphere at 37°C. All cell lines are frequently tested for mycoplasma contamination.

### Herpes simplex virus 1 stocks and infections

HCT116 cells were infected with HSV1(17+) or HSV1(17+)Lox-_pMCMV_GFP at a MOI of 0.04^32,33^. 4 days post infection (dpi), the supernatant was harvested, transferred to 50 ml tubes and centrifuged at 3000×g and 4°C for 5 min to remove cell debris. Virus was enriched from the remaining supernatant by size excluding filtration using Centricon Plus-70 (Merck) with a threshold of 100 kDa. Beforehand, Centricons had been washed with H_2_O, disinfected with 70% ethanol and again washed with H_2_O. The corresponding centrifugation steps were carried out at 1000×g and 4°C for 15 min. 60 ml of virus-containing supernatant were transferred into single Centricons and subsequent centrifugation was performed at 1000×g and 4°C for 60 min. Enriched virus stocks were eluted in the provided concentrate recovery tubes by a 2 min centrifugation step at 1000×g and 4°C. Vital titers were determined by plaque assays on VeroE6 cells.

### Adenovirus C5 stocks and infections

A replication-competent HAdV-C5 delta E3 virus, encoding a CMV promoter-driven eGFP expression cassette, was further used in this study^34^. The virus was propagated and titrated in HEK293 cells. Virus titer was determined by immunofluorescence staining of the adenoviral DNA binding protein E2A^35^. A549 cells were infected with HAdV-C5 delta E3 virus at a MOI of 0.5 in non-supplemented DMEM. After incubation for 1 h at 37°C, the infection was stopped by replacing virus containing medium with fresh DMEM medium with supplements. Degron-tagged HCT116 cells were infected with HSV-1 strain 17 at a MOI of 0.5. After an incubation period of 30 min at 37°C and 5% CO_2_, medium was discarded and conditioned medium added. At defined times post infection, samples were harvested.

### RNA interference

HCT116 or A549 cells were seeded out in a 12-well format at a density of 4x 10^4^ or 3x 10^3^ cells per well, respectively. Cells were transfected with respecting siRNAs using RNAiMAX (Thermo Fisher). In short, 671 μl OptiMEM (Thermo Fisher) with 4.5 μl siRNA (20 μM), and 648 μl OptiMEM with 27 μl RNAiMAX were prepared. The mixtures were combined and had been incubated for 10 min at room temperature. The final amount of siRNA per well was 8.34 pmol. Catalog numbers of siRNAs used in this study are listed in Table 2. 48 hours post transfection (hpt) supplemented media was added. The following day infections were performed.

**Table 2:** List of used siRNAs from Horizon Discovery/Dharmacon.

| Catalog Number | Product Name |
| --- | --- |
| J-012286-05-0002 | ON-TARGETplus Human SLBP (7884) siRNA - Individual, 2 nmol |
| J-011548-05-0002 | ON-TARGETplus Human FUBP1 (8880) siRNA - Individual, 2 nmol |
| J-006455-06-0002 | ON-TARGETplus Human SFPQ (6421) siRNA - Individual, 2 nmol |
| J-016234-05-0002 | ON-TARGETplus Human SUPT5H (6829) siRNA - Individual, 2 nmol |
| J-005150-05-0002 | ON-TARGETplus Human SAFB (6294) siRNA - Individual, 2 nmol |
| D-001810-01-05 | ON-TARGETplus Non-targeting siRNA #1, 5 nmol |

### RNA/DNA isolation

For isolation of cellular RNA and DNA, cells were thoroughly resuspended in 300 μl Trizol (ThermoFisher). If both RNA and DNA were extracted from Trizol, one third was processed using the DNA Trizol kit (Zymo). The remaining sample or otherwise the entire 300 μl were used as input for the Direct-zol RNA miniprep kit (Zymo) to digest DNA and extract RNA by following the manufacturer’s instructions. RNA integrity was confirmed by running an RNA ScreenTape Analysis (Agilent Technologies) on a TapeStation and concentrations were determined by NanoDrop measurements.

### Reverse transcription and qPCR

500 ng of extracted RNA from each sample was reverse transcribed with the SuperScript™ III Reverse Transcriptase (ThermoFisher) according to the manufacturer’s recommendations using RNaseOUT ™ Recombinant RNase Inhibitor (ThermoFisher) and 100 ng random primers (ThermoFisher). The resulting cDNA was diluted 1:20 with DNA-/RNA-free water. Per qPCR reaction, 3.75 μl of the diluted cDNA were amplified using the GoTaq qPCR Kit (Promega) and 1 μM each of the previously described primers, shown in Table 3^36^. For normalization purposes, raw UL54 RNA values were normalized using GAPDH, and raw UL54 DNA values using the BBC3as primer pair which targets the human genome.

**Table 3.**
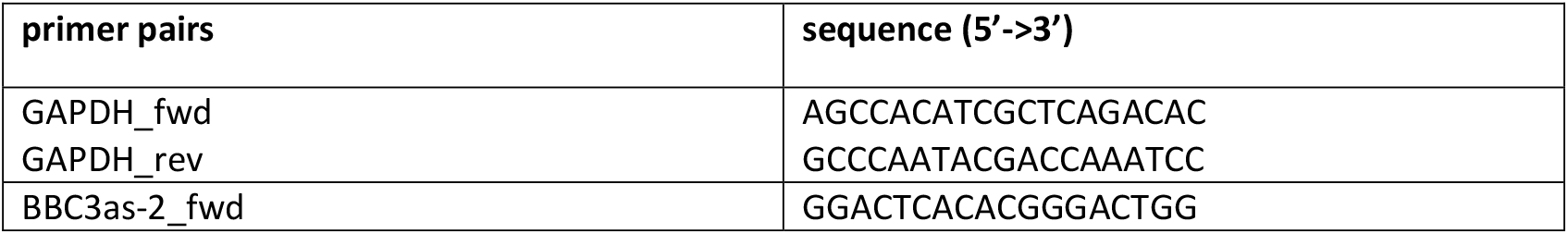
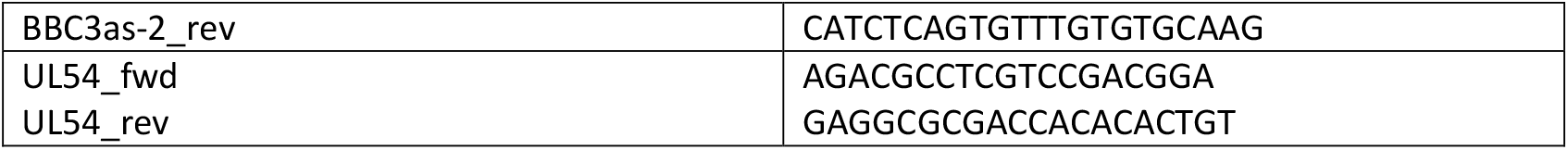
primers.

### Preparation of virus stocks for co-localization experiments

To produce EdC-labeled viruses, HaCaT cells were seeded in 10× 15 cm dishes (Day 0). On the following day, cells were washed once with PBS and infected with HSV-1 strain 17 (1.2×107 PFU/ml) at a MOI of 0.01, using 2 ml inoculum per dish. At 1 h post-infection (hpi), 1.5 μM Ethynyl-dCTP (EdC) (ThermoFisher) was added to four plates per virus (in 18 ml medium per plate). Controls (one plate per virus) received an equivalent volume of DMSO. On Days 2 and 3, 1.5 μM EdC or the equivalent amount of DMSO was added to the respective plates. On Day 4, virus-containing supernatants were harvested, and pre-cleared by centrifugation (500×g, 5 min, 4°C). Supernatants were filtered through a 0.45 μm membrane, and concentrated by centrifugation (14.000×g, 1.5 h, 4°C) over a 5% OptiPrep cushion. The virus pellet was resuspended in 500 μl PBS. Viral titers were determined by performing plaque assays on Vero cells.

### Western blot analysis

For protein analysis, cells were resuspended in RIPA lysis buffer (50 mM Tris-HCl [pH 8,0], 150 mM NaCl, 5 mM EDTA, 1% [vol/vol] Nonident P-40, 0.1% [wt/vol] SDS, 0.5% [wt/vol] sodium deoxycholate) supplemented with a fresh protease inhibitor cocktail containing 0.2 mM PMSF, 1 mg/mL pepstatin A, 5 mg/mL aprotinin, and 20 mg/mL leupeptin. After 30 min incubation on ice, the lysates were sonicated and cell debris was pelleted at 14.800 g at 4°C for 3 min. Cell lysates were boiled for 3 min at 95°C in 5x Laemmli buffer. Proteins were separated by SDS-PAGE, transferred to nitrocellulose blotting membranes (0.45 μm) and visualized by immunoblotting.

Primary antibodies specific for HAdV proteins used in this study included mouse monoclonal antibody (Mab) mouse B6-8 (anti-E2A), MAb mouse AC-15 (anti-β-actin; Sigma-Aldrich)^35^. Primary antibodies specific for cellular expressed proteins included HNRNPU (abcam; Cat# ab10297), SAF-B (Santa Cruz Biotechnology; Cat# sc-135618), SFPQ (abcam; Cat# ab38148), SLBP (abcam; Cat# ab181972), SUPT5H (abcam; Cat# ab126592), FUBP1 (abcam; Cat# ab181111), GAPDH (Sigma-Aldrich; Cat# G8795).

Secondary antibodies conjugated to horseradish peroxidase (HRP) for detection of proteins by immunoblotting were anti-rabbit IgG, anti-mouse IgG, and anti-rat IgG (Jackson/Dianova or Dako) used in a dilution of 1:10.000.

Degron-tagged HCT116 cells subjected to RNA extraction and subsequent RT-qPCR were seeded at a density of 8x 10^6^ cells per well in a 12-well plate. Before the infection, cells were treated with 500 μM indole-3-acetic acid (IAA) (Sigma-Aldrich) for 4h or 22h to induce protein depletion or remained untreated, respectively. The infection was carried out with HSV1(17+)Lox-_pMCMV_GFP at a MOI of 1. After an incubation period of 30 min at 37°C and 5% CO_2_, medium was discarded and conditioned medium added. At 2hpi and 4hpi, the medium was discarded and cells were lysed in 300 μl Trizol (ThermoFisher).

### Fixation of infected cells and fluorescence microscopy

For the purpose of imaging, immunofluorescently labeled RBPs and HSV-1EdC-labeled genomes, parental HCT116 cells and degron-tagged cell lines were seeded at 50.000 cells per well in 8-well plates (ibidi). Cells were pretreated with 500 μM IAA (Sigma-Aldrich;) 4 hours before infection to induce protein depletion where applicable. Cells were infected with HSV-1EdC or HSV-1DMSO at an MOI of 10. At 3 hpi, cells were washed with PBS and fixed with 4% PFA for 15 minutes at room temperature (RT). Following fixation, samples were washed three times with PBS, permeabilized with 0.1% Triton X-100 for 20 minutes, and blocked with 3% BSA for 15 minutes at RT. Following two additional PBS washes, the click chemistry reaction was performed with the Click-iT Plus Alexa Fluor 488 Picolyl Azide Toolkit according to the manufacturer’s protocol (ThermoFisher). After the click reaction, cells were washed once in 3% BSA and twice with PBS.

Quantitative image analysis was performed in ImageJ Fiji (Schindelin et al., 2012) using the plugin Trackmate v7.13.2 (Ershov et al., 2022) to identify viral genomes, applying a detection threshold of 750. The median intensity of each protein was then measured. To establish protein-specific intensity thresholds, the median intensity was determined at three randomly selected nuclear positions. The threshold was set at 1,000 for SUPT5H and 2,500 for SFPQ, since SFPQ exhibited a higher baseline intensity. Colocalization events were defined as signals exceeding the respective thresholds, and the proportion of colocalized spots relative to DNA signals was calculated. A total of 95 and 118 genomes were analyzed for SUPT5H and SFPQ, respectively.

For immunostaining, cells were incubated with an anti-V5 primary antibody (1:500) (Cell Signaling) for 1 hour at RT, washed three times with PBS, and incubated with a goat anti-rabbit secondary antibody (1:1000) (ThermoFisher). Nuclei were counterstained with Hoechst (1:1000) (ThermoFisher). Samples were imaged using a Spinning Disk microscope (Nikon).

To evaluate viral transcription/replication, the signal of GFP expressing HSV1 or HAdV was imaged and measured, respectively.

## Results

### siRNA-mediated knockdown of RBPs involved in transcriptional regulation impairs HSV-1 and HAdV gene expression while reducing cell growth

To investigate the impact of cellular RBPs on early HSV-1 transcription, we initially focused on five candidates with known nuclear localization. These five RBPs were selected based on previous reports, such as an HSV-1 siRNA knockdown screen.^37^

The first protein of interest, Far Upstream Element Binding Protein 1 (FUBP1), was mainly described as a transcriptional activator of c-myc expression, which in turn acts as a transcription factor for cell growth and proliferation^38^. Mechanistically, FUBP1 interacts with transcription factor II H (TFIIH), an essential component of pre-initiation complexes, to facilitate promoter escape of RNA polymerase II (RNAPII) by stimulating TFIIHs helicase activity^38,39^. While knockdown of FUBP1 was already described to reduce HAdV replication, its impact on HSV-1 replication is unknown^40^.

At the beginning of infection, HSV-1 genomes brought into the cell by virus particles are free from histones but quickly get chromatinized after their entry into the nucleus^41–43^. Similar to the host, viral transcription depends on epigenetic regulation allowing or restricting access of the transcriptional machinery to the genome. Whereas cellular genes are regulated individually due to different chromatin states, the epigenetic regulation of HSV-1 DNA occurs on a genome-wide level with transcription factors controlling the fine-tuned activation of separate viral genes^44^. Histone modifications on the entire HSV-1 genome could therefore determine globally silenced or permissive chromatin states and thus dictate whether cells undergo lytic or latent infection^44^. Since the Stem-Loop Histone mRNA Binding Protein (SLBP) is required for processing histone mRNAs, we wondered whether its depletion would affect viral transcription competence^45,46^.

Another RBP of interest, Splicing Factor Proline And Glutamine Rich (SFPQ), is described to have multiple functions that depending on the virus under investigation can be pro- or antiviral^47–51^. Besides its role in alternative splicing, SFPQ acts as a transcriptional repressor and is a component of paraspeckles, which are nuclear entities sequestering RNA molecules for RNA editing and are further involved in transcriptional regulation of HSV-1 genes ^26,28,52–54^.

Spt5, encoded by the SPT5 Homolog, DSIF Elongation Factor Subunit (SUPT5H) gene, is part of the DSIF (DRB-sensitivity inducing factor) complex that associates with RNA polymerase II and affects promoter-proximal pausing and elongation^55^. Moreover, SUPT5H was previously shown to be particularly important for the transcription of G+C-rich genes^56^. As those are highly prevalent in the HSV-1 genome, SUPT5H is likely relevant for viral transcription as well, possibly through an interaction with the viral protein Icp22.

Finally, the Scaffold attachment factor B (SAFB) was described to link transcription to splicing and thus represents an interesting candidate to study in the context of viral infection^57^.

To assess the role of these five RBPs in early viral transcription, we depleted their mRNAs in A549 cells and HCT116 cells using siRNAs for 72 h, resulting in a strong reduction in protein levels (Supplementary Figure 1A). At this time point, we infected cells with 0.5 MOI of HSV-1, and quantified the immediate early transcript UL54 by RT-qPCR at 2 hpi (hours post infection) (Figure 1A). Depletion of SFPQ and SUPT5H resulted in a strong reduction in viral RNA levels, whereas the other three candidates showed no substantial effect. However, vDNA levels remained largely unchanged, indicating that viral entry and genome delivery were not impaired by the knockdowns (Figure 1B).

**Figure 1:**
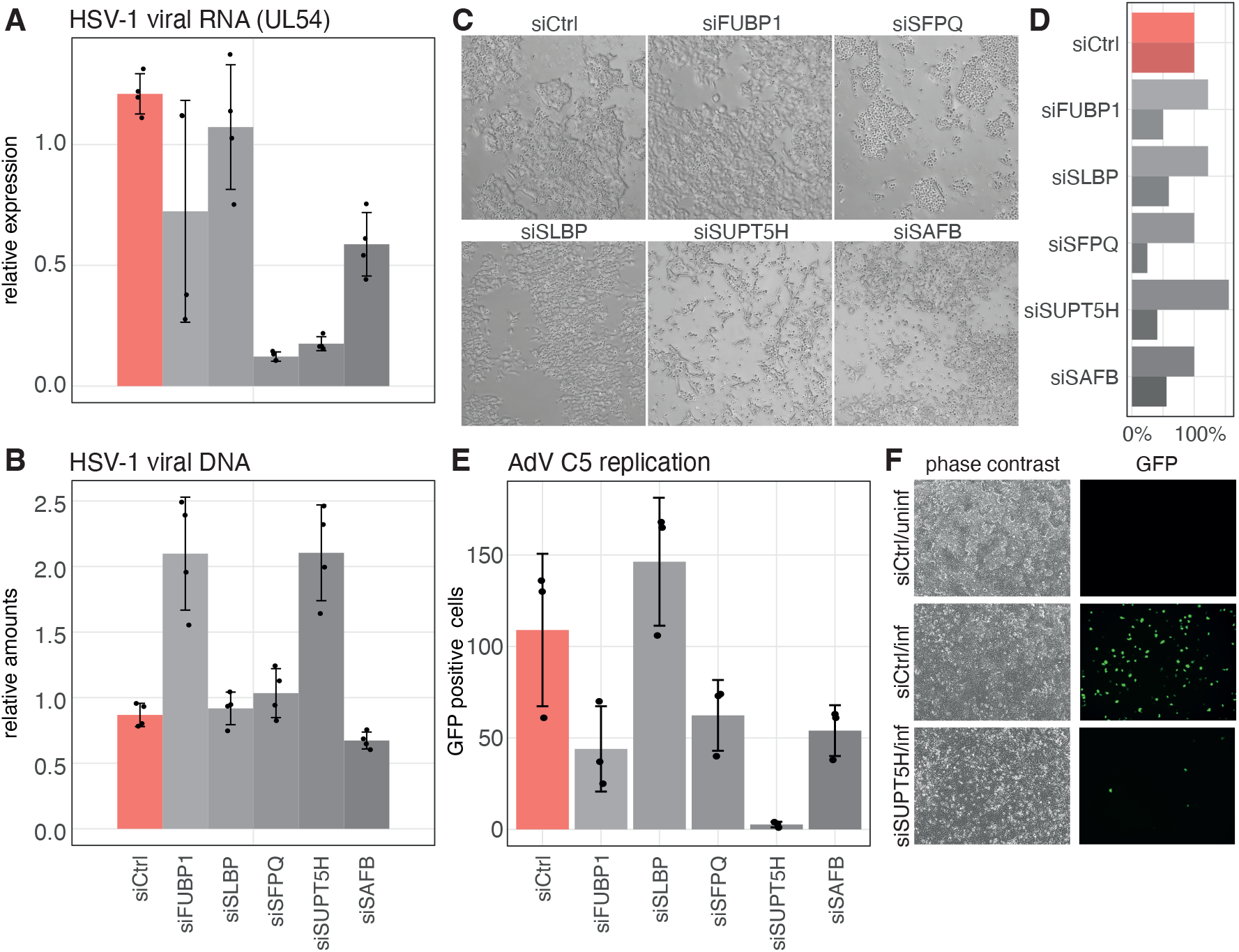
RNA interference to study influence of RNA binding proteins on herpes simplex virus 1 and adenovirus C5 infection. **A**, HCT116 cells were transfected with the indicated siRNAs. At 72 h post transfection, cells were infected with an MOI of 0.5. Further 2 h later, intracellular RNA was collected, and viral RNA (UL54 transcript) quantified by RT-qPCR. **B**, as in A, but for intracellular viral DNA quantified by qPCR. **C**, phase contrast microscopy of the cells used for Fig. 1A at 72 h after transfection of the indicated siRNA. **D**, for the same experiment as in C, the barplot shows for each treatment indicated on the left, relative to siCtrl, the number of seeded cells (upper, brighter bar) and cell count 72 h after RNAi (lower, darker bar). **E**, A549 cells were transfected with the indicated siRNAs. At 48 hpost transfection, cells were infected with 0.5 MOI GFP-expressing adenovirus C5, and again 24 hlater GFP signal from the virus was quantified. Shown are median and standard deviation from three measurements. **F**, representative microscopy images for, from top siCtrl/not infected, siCtrl/infected, siSUPT5H/infected.

Microscopic inspection of cells after 72 h of RNAi revealed altered cell morphology in some conditions compared to the control siRNA treatment (Figure 1C). In additon, cell counts at seeding and harvesting indicated that knockdowns of SFPQ and SUPT5H considerably impaired cell growth (Figure 1D).

For our comparative approach with human adenovirus C5 (HAdV C5), we carried out the RNAi experiment in A549 cells using a GFP-expressing adenovirus. Cells were infected 48 h post siRNA transfection, and viral replication was assessed 24 h later by counting GFP-positive cells (Supplementary Figure 1B). Similar to HSV-1, the knockdown of SUPT5H strongly impaired viral replication (Figure 5E+F). However, the knockdown of SFPQ did not produce a strong effect, suggesting that parallel experiments with different DNA viruses can help to distinguish general host factors essential for nuclear-replicating DNA viruses from those specific for HSV-1.

Taken together, while comparative viral studies provide valuable insight, the prolonged RNAi treatment required for robust protein depletion significantly affects cell morphology and growth. These secondary effects may well complicate functional analyses of essential regulatory RBPs, raising the possibility that observed differences in viral transcription and replication result from secondary effects of RNAi rather than direct involvement of the proteins of interest.

### Generation of AID-tagged cell lines and validation of RBP depletion

To more precisely assess the direct consequences of specific RBP depletion on viral transcription, we utilized an auxin-inducible degron (AID) system. In HCT116 cells stably expressing the F-box protein *At*AFB2 (HCT116 AFB2 cells), we endogenously tagged SFPQ and SUPT5H with a miniIAA7 tag (AID-tag) (Figure 2A). Upon addition of indole-3-acetic acid (auxin), this tag allows for the recruitment of the exogenously expressed plant *At*AFB2 receptor that interacts with cellular components of an E3-ubiquitin ligase complex. As a result, the tagged protein undergoes polyubiquitinylation and is rapidly degraded by the proteasome. To validate the depletion of SUPT5H and SFPQ, we treated AID-degron tagged HCT116 AFB2 cells with auxin and monitored protein levels over a 24 h time course by Western blot analysis. We observed a significant reduction in SUPT5H and SFPQ protein levels within 2 h and 4 h after the addition of auxin, respectively confirming the functionality of our genome-edited HCT116 AFB2 cells (Figure 2B).

**Figure 2:**
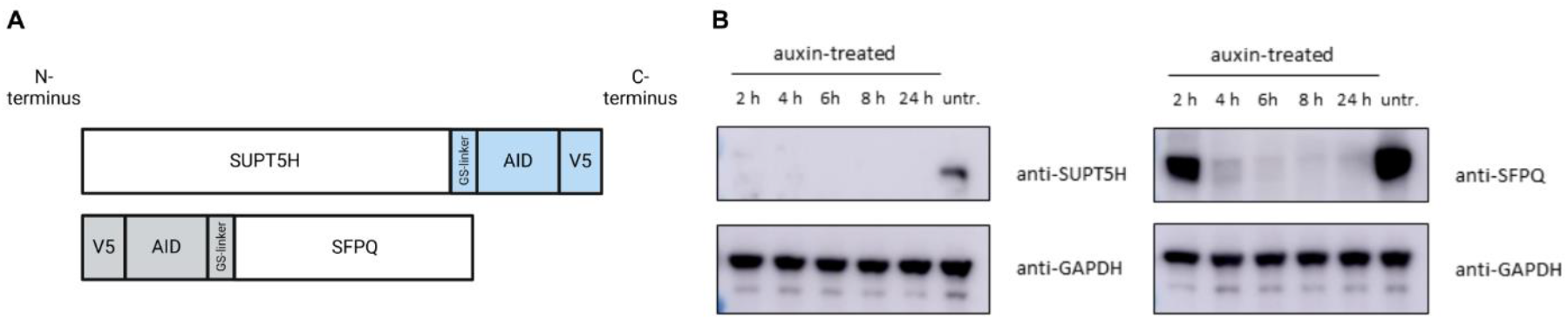
Auxin-inducible degradation of SFPQ and SUPT5H. **A**, host RNA-interacting proteins SFPQ and SUPT5H were tagged at N-(grey) or C-terminus (blue) with a Glycine/Serine-linker, a miniIAA7 degron, and a V5-tag. Schemes of the modified proteins are depicted. Created in BioRender. Landthaler, M. (2025) https://BioRender.com/w18y704 **B**, HCT116 AFB2 cells with miniIAA7 (AID)-tagged proteins were treated with 500 μM indole-3-acetic acid (auxin) or left untreated (untr.). At indicated times after treatment, cells were lysed and a Western blot analysis was performed with 20 μg of each protein sample loaded per lane. Blots were probed with protein-specific antibodies indicated on the right.

In summary, by using the AID system, we achieved rapid and robust depletion of SUPT5H and SFPQ within 4 h, providing a considerably faster alternative to RNAi knockdowns.

### Reduction of early viral transcription in SFPQ-degron cells depends on the timespan of depletion of infection

To determine whether RBP depletion using the AID system affects early viral transcription, we infected cells 4h (SUPT5H and SFPQ) or 22h (SFPQ) after auxin addition (Figure 3A). At 2 and 4 hpi, we extracted total RNA from the cells, and quantified viral transcription by RT-qPCR of the early gene UL54. For SUPT5H, depletion of the protein for 4h prior to the infection resulted in a strong reduction of vRNA at both 2 hpi and 4 hpi (Figure 3B). This can be expected since SUPT5H is a component of the RNAPII machinery. Alternatively, the deletion of SUPT5H had been reported to destabilize a core component of RNAPII, namely RPB1^58^.

**Figure 3:**
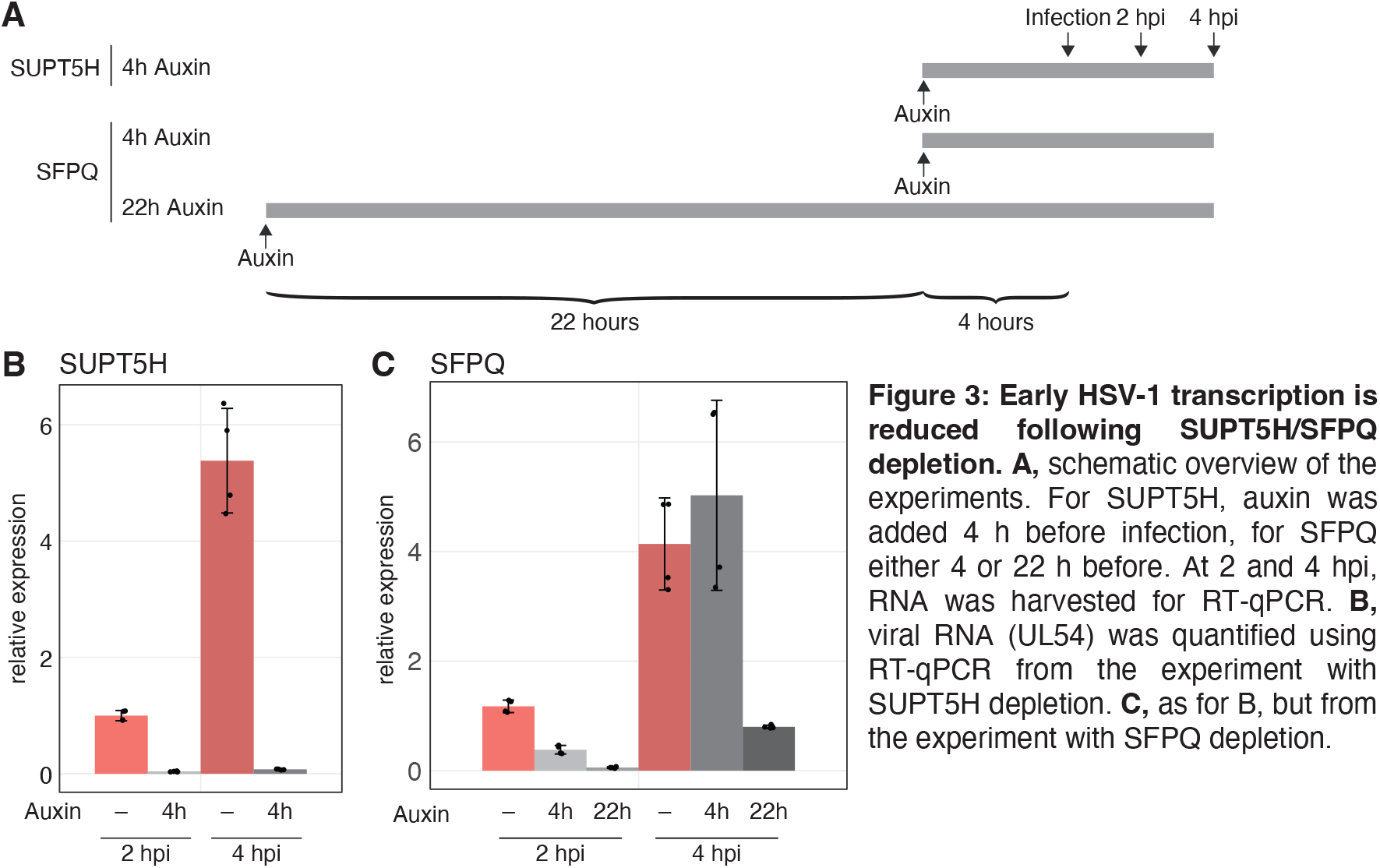
Early HSV-1 transcription is reduced following SUPT5H/SFPQ depletion. **A**, schematic overview of the experiments. For SUPT5H, auxin was added 4 h before infection, for SFPQ either 4 or 22 h before. At 2 and 4 hpi, RNA was harvested for RT-qPCR. **B**, viral RNA (UL54) was quantified using RT-qPCR from the experiment with SUPT5H depletion. **C**, as for B, but from the experiment with SFPQ depletion.

In contrast, SFPQ depletion for 4h before infection decreased vRNA levels by about 60% at 2hpi. However, at 4hpi viral transcription had recovered, showing no difference from untreated cells (Figure 3C). Interestingly, when SFPQ was depleted for 22 hours before infection, the reduction in vRNA effect at 2hpi was more pronounced compared to the shorter depletion, and was also persistent at 4hpi (Figure 3C). This suggests that prolonged depletion of SFPQ might lead to secondary effects such as disintegration of paraspeckles since SFPQ is critical for the structural integrity of these subnuclear bodies^59^. Thus, the observed reduction in viral transcription might stem from broader nuclear structural alterations rather than a direct role of SFPQ in viral transcription or RNA processing. The findings highlight the advantage of the AID system in distinguishing primary effects of RBPs depletion from secondary consequences, enabling a more precise dissection of host factors involved in viral transcription.

### The transcription elongation factor SUPT5H, but not SFPQ, co-localizes with HSV-1 genomes in early infection

Given that both, SUPT5H and SFPQ, affect viral mRNA levels, we sought to determine whether they act directly on viral genomes to regulate transcription or modulate HSV-1 gene expression through alternative mechanisms. Since protein co-localization with DNA is a strong indicator of transcription-regulated functions, we investigated whether SUPT5H and SFPQ are in close-proximity to viral genomes. To this end, SUPT5H- and SFPQ-degron cells were infected with HSV-1 either containing EdC-labeled or unlabeled (DMSO control) genomes. At 3 hpi, cells were fixed and stained with a V5-specific antibody to detect SFPQ or SUPT5H, while viral DNA was fluorescently labeled using click-chemistry.

We observed distinct SUPT5H foci at sites of fluorescently labeled HSV-1 genomes, which were absent upon SUPT5H depletion (Figure 2A and Supplementary Figure 2A), indicating its accumulation at viral DNA. In contrast, SFPQ did not show increased signal intensities at HSV-1 genomic sites (Figure 2B). To confirm our visional impression, we quantified the relative number of viral DNA foci at locations of protein enriched foci. Indeed, 89% (SD= 1,45%) of vDVA foci occurred at spots with accumulated SUPT5H signal, whereas the relative number of SFPQ and vDNA co-localization were markedly lower (16%; SD=13,3%) (Supplementary Figure 3).

While the association of SUPT5H with viral DNA aligns with the dependence of HSV-1 on the host transcriptional machinery and the role of SUPT5H as a transcription elongation factor, we found no evidence of SFPQ co-localization with HSV-1 genomes. Thus, SFPQ likely influences viral gene expression through a mechanism independent of direct interaction with viral DNA.

## Discussion

In this study, we initially focused on 5 RNA-interacting proteins (FUBP1, SLBP, SFPQ, SUPT5H and SAF-B) that reside in the nucleus. Although their involvement in host transcriptional regulation has been reported already, mechanistic insights are still elusive or their contribution to viral transcription is largely unknown. We conducted protein depletion experiments and measured the expression of the immediate-early gene UL54 in order to examine the consequences for HSV-1 transcription. Of our 5 candidates, siRNA-mediated knockdown of SUPT5H or SFPQ resulted in the most pronounced decrease of UL54 mRNA. To gain a better understanding if these RNA interacting proteins are exclusively important for HSV-1 gene expression or have broader implications for the replication of dsDNA viruses, we extended our experiment to adenovirus C5. We observed that the knockdown of SUPT5H drastically reduces the growth of adenovirus C5. In contrast, depletion of SFPQ did not result in prominent changes, which confirms a previous finding that knockdown of SFPQ using RNAi does not affect adenovirus titer^51^.

SUPT5H positively regulates several events during RNAPII-mediated transcription beginning with promoter-proximal pause release and followed by elongation and termination^58,60^. Due to SUPT5Hs central role in global host transcription, we anticipated severe consequences for viral gene expression upon its depletion, which holds true. Our results thus suggest that SUPT5H is not only required for the host but also for HSV-1 and adenovirus C5 transcription, which is further in line with our evidence showing its co-localization with HSV-1 genomes. It has been reported that depletion of SUPT5H results in the degradation of RPB1, a core subunit of RNAPII, which could mechanistically explain our results^58^. The degradation happens during early elongation and is controlled by CDK9^58^. Similar to a previous study, inhibition of CDK9 and subsequent induction of SUPT5H depletion at the start of infection could allow us to analyze SUPT5H’s importance for elongation and termination of viral genes without being stopped by RPB1 degradation^58^.

In contrast to SUPT5H, SFPQ was not found in co-localization with HSV-1 genomes in our study, indicating a function that is not exerted on viral DNA during early times of infection. SFPQ is pivotal for the formation of paraspeckles, nuclear entities consisting of NEAT1_2 lncRNA, PSPC1, NONO, and SFPQ, which were shown to be induced upon different cellular stress including HSV-1 infection^52,59,61–63^

Despite RNA processing, nuclear retention of RNAs and transcriptional regulation have all been attributed to paraspeckle formation, their precise function especially during infection is not fully understood^28,64,65^. Considering that depletion of SFPQ disintegrates paraspeckles, it might affect HSV-1 transcription even though the underlying mechanism needs to be clarified^59^.

On the other hand, SFPQ acts as a transcriptional repressor that is relieved during infection, likely redistributes to paraspeckles and thereby allows expression of genes involved in the immune response, as it was described already for IL8 and RELA^52,66^. This mechanism could establish an antiviral state of cells that are being primed before the HSV-1 infection even started and could in part explain why cells that had been subjected to siRNA-mediated knockdown for multiple days or auxin-inducible degradation of SFPQ over an extended timespan (22 h) showed a stronger decrease in UL54 RNA levels compared to a faster depletion strategy (4 h).

Furthermore, this observation exemplifies the advantage of the AID system over RNAi. Due to relatively long half-lives of proteins, RNAi experiments often suffer from long incubation periods of siRNA-transfected cells in order to achieve target depletion. In addition, studies on the function of essential genes by deletion or RNAi are inherently restricted by the loss of cell viability. As demonstrated in our approach for SFPQ, the duration of protein depletion matters since it influences the observed effect size. While secondary effects of protein depletion can never be fully ruled out, the AID system minimizes the period of time for these effects to occur. Owing to its temporal flexibility, it further helps to dissect immediate and belated consequences of protein depletion. Currently, only a few studies in the field of virology take advantage of auxin-inducible degradation. However, we are convinced by its potential to foster our understanding of how RNA-interacting proteins and other essential host factors contribute to viral transcription and replication.

## Supporting information

Supplementary Figure S1 Part1

Supplementary Figure S1 Part2

## Acknowledgments

We thank Ouidad Benlasfer and Nouhad Benlasfer for technical assistance, as well as members of the Landthaler lab for discussions and comments. This work was supported by the Deutsche Forschungsgemeinschaft (DFG; German Research Foundation) in the framework of the Research Unit FOR5200 DEEP-DV (443644894) project BO 4158/5-1, SCHR 1479/5-1 and LA 2941/18-1 to J.B., S.S. and M.L., respectively.

---

**Figure 4:**
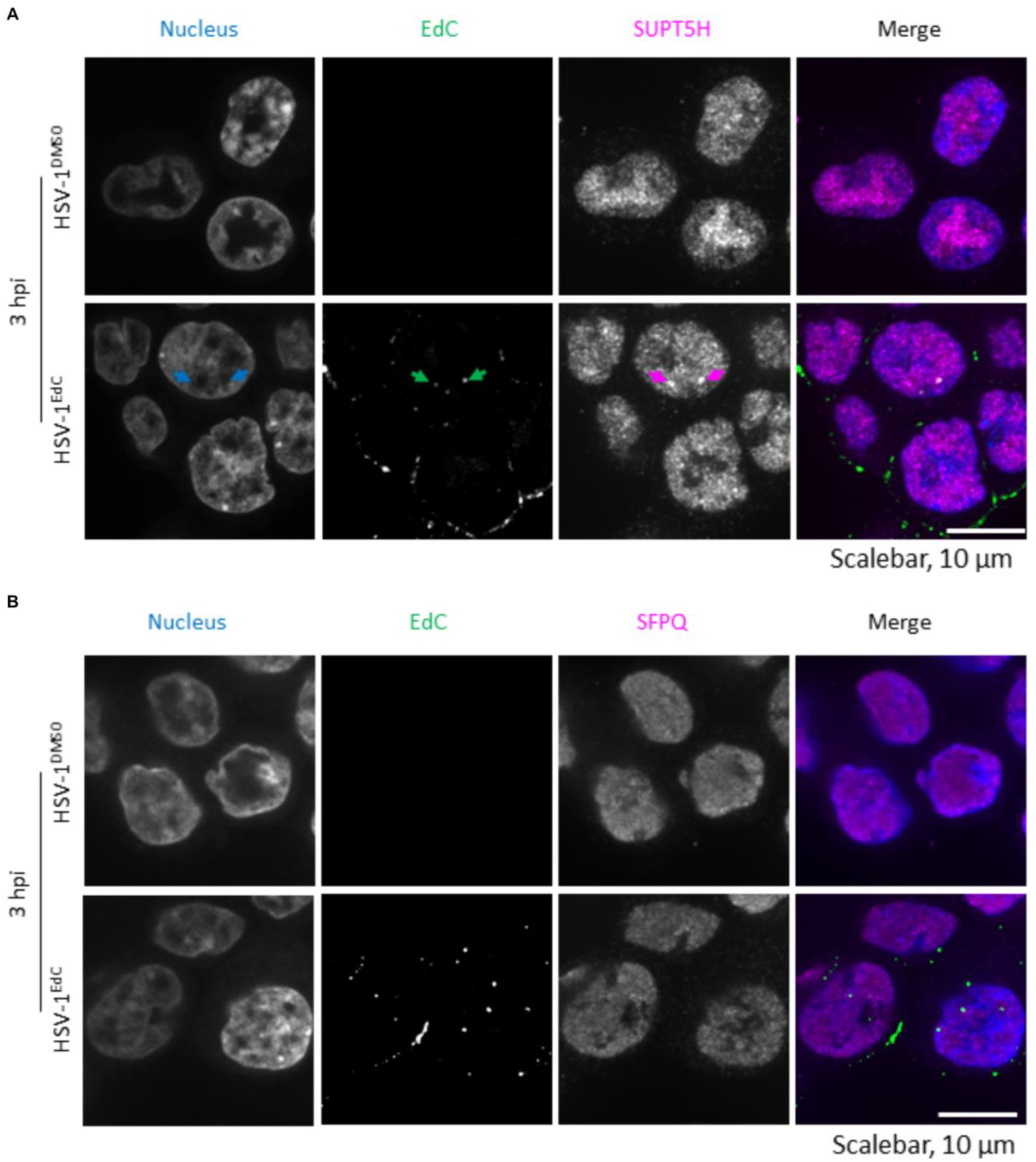
SUPT5H co-localizes with viral genomes early in HSV-1 infection. **A**, SUPT5H-degron cells were either infected with HSV-1 carrying unlabeled (DMSO) or EdC-labeled DNA. At 3 hpi, all cells were fixed, their nuclei were stained with Hoechst, viral DNA was fluorescently stained by a click reaction, SUPT5H was immunostained with a V5-antibody and images were taken. Arrows indicate a signal of viral DNA. **B**, as in A, but for SFPQ-degron cells.

