## Supplementary Figure S1 Part1 for "Dissecting the role of RNA-binding proteins in early herpes simplex virus 1 transcription using acute protein depletion"

**A**

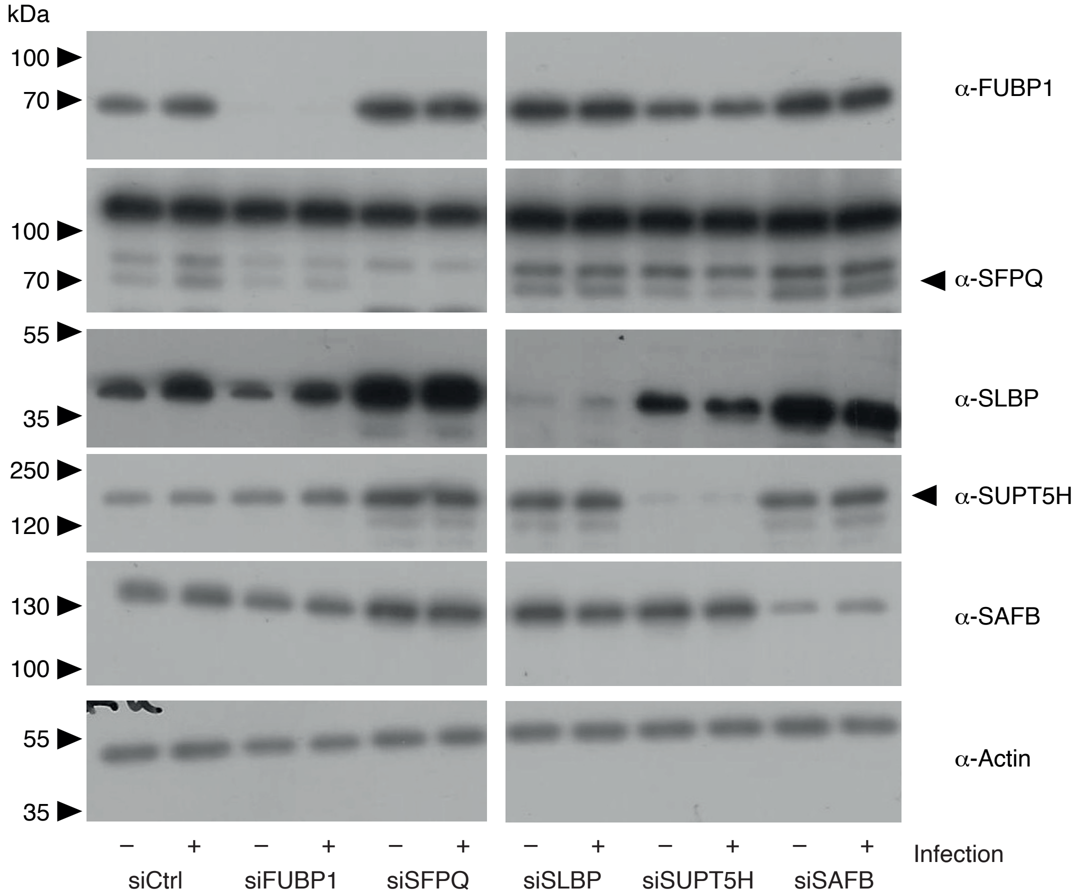


**Supplementary figure 1: A,** Western blot analysis of protein depletion by indicated. The employed antibodies are indicated on the right. Theoretical weights are as follwing: FUBP1 – 70kDa, SFPQ – 100 kDa, SLBP – 40 kDa, SUPT5H – 150 kDa, SAFB – 150 kDa. **B,** A549 cells were infected with a GFP expressing AdV C5, and visualized in bright field (left column) and the GFP channel (middle column). Top row, uninfected cells, other rows as indicated.
