## Supplementary figures and images for "Dissecting the role of RNA-binding proteins in early herpes simplex virus 1 transcription using acute protein depletion"

### Supplementary Figure S1 Part2

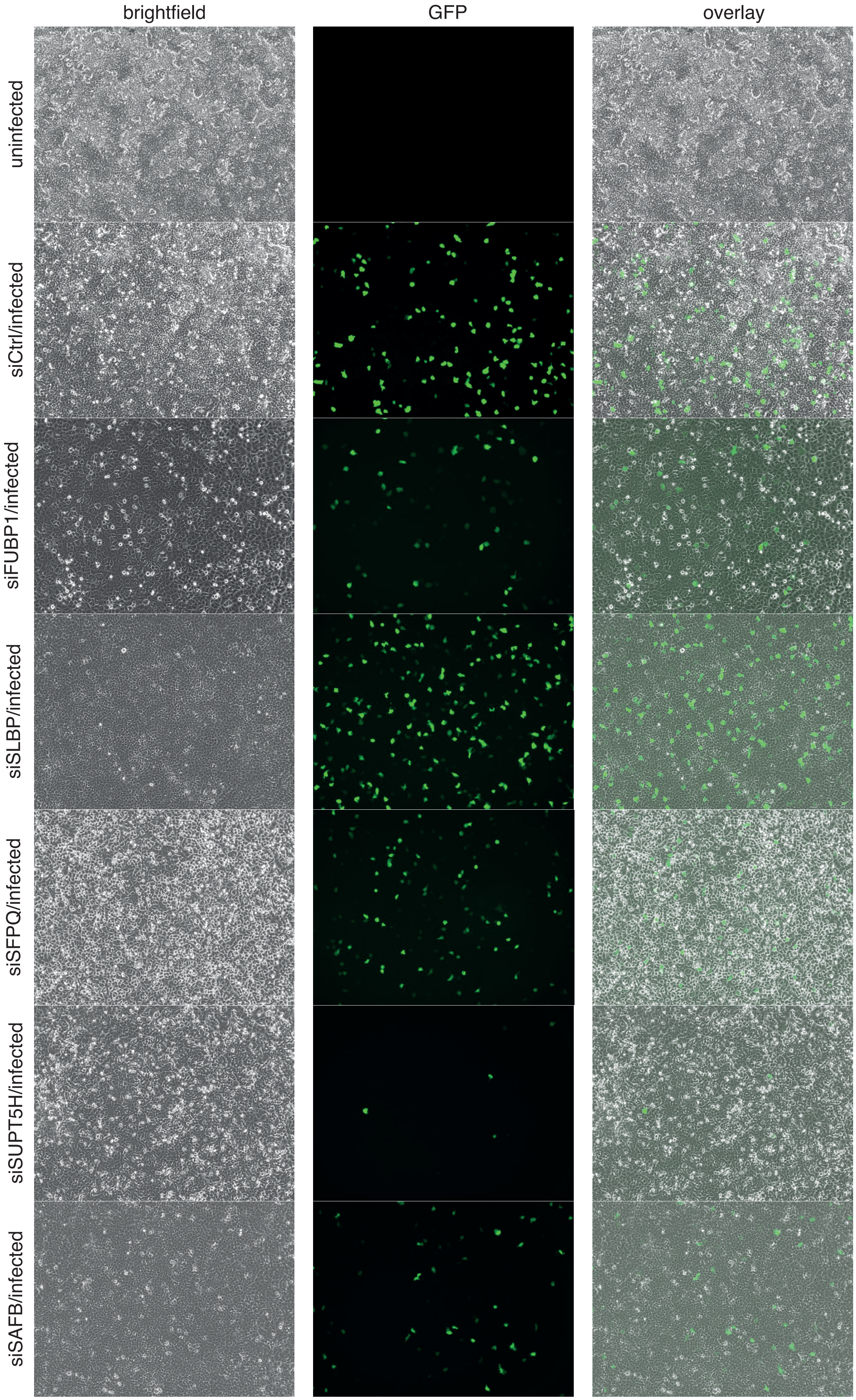
