## Supplementary Figure S2 for "Dissecting the role of RNA-binding proteins in early herpes simplex virus 1 transcription using acute protein depletion"

A

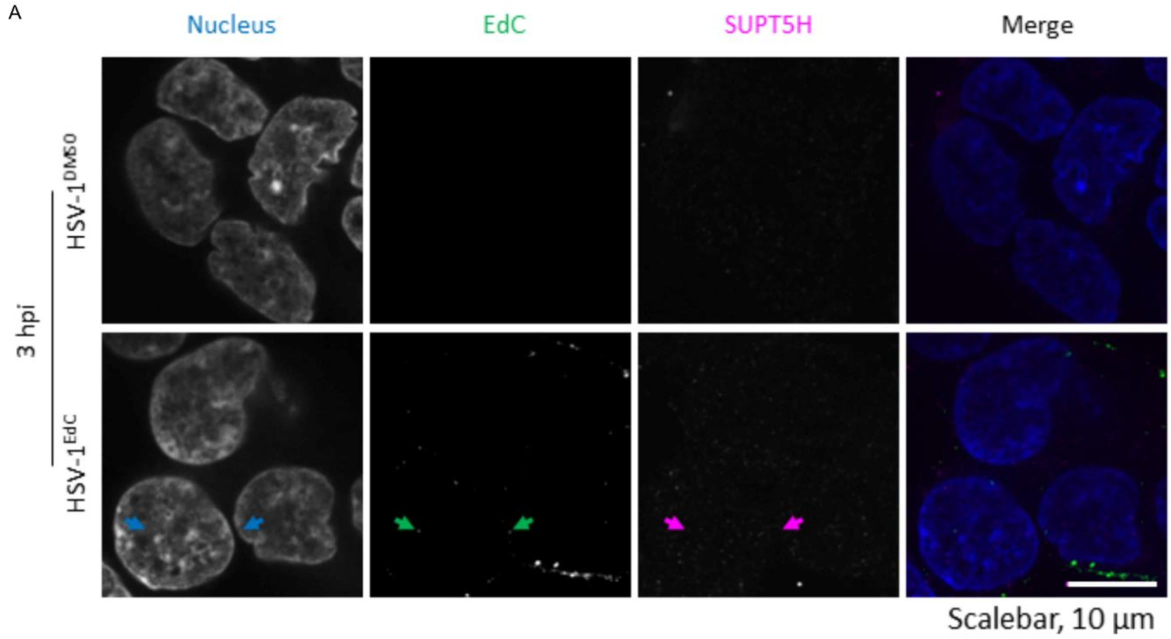


B

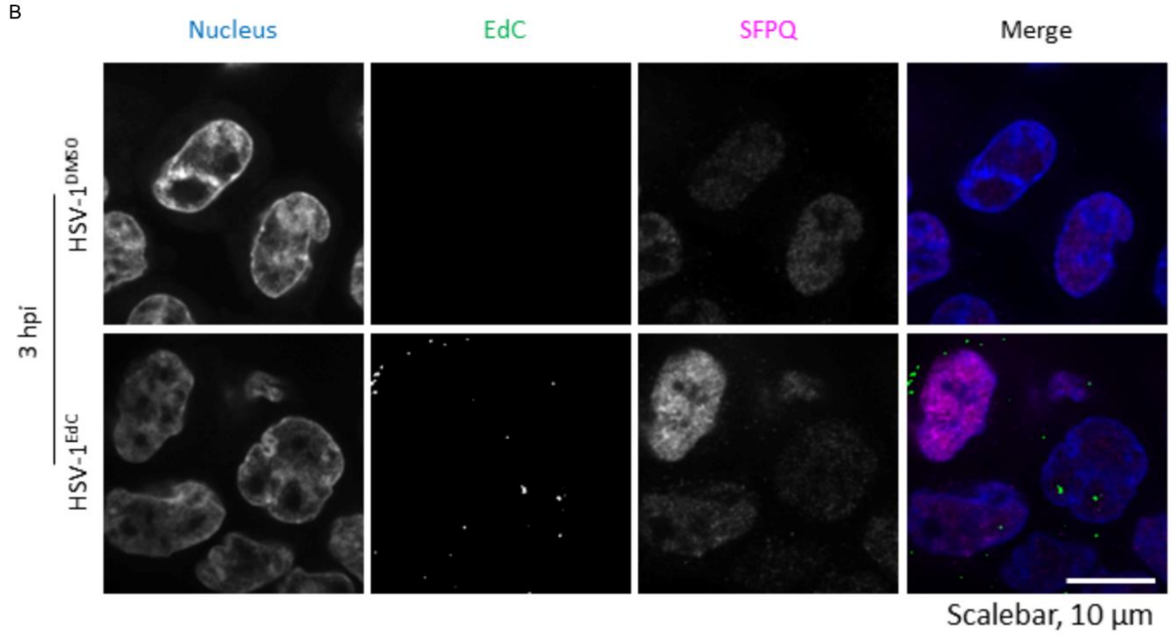


**Supplementary Figure 2:** Auxin-induced depletion of SUPT5H and SFPQ serve as negative control. **A**, SUPT5H-degron cells were treated with 500 µM indole-3-acetic acid (auxin), before they were either infected with HSV-1 carrying unlabeled (DMSO) or EdC-labeled DNA. At 3 hpi, all cells were fixed, their nuclei were stained with Hoechst, viral DNA was fluorescently stained by a click reaction, SUPT5H was immunostained with a V5-antibody and images were taken. Arrows indicate a signal of viral DNA. **B**, as in A, but for SFPQ-degron cells.
