## Supplementary Figure S3 for "Dissecting the role of RNA-binding proteins in early herpes simplex virus 1 transcription using acute protein depletion"

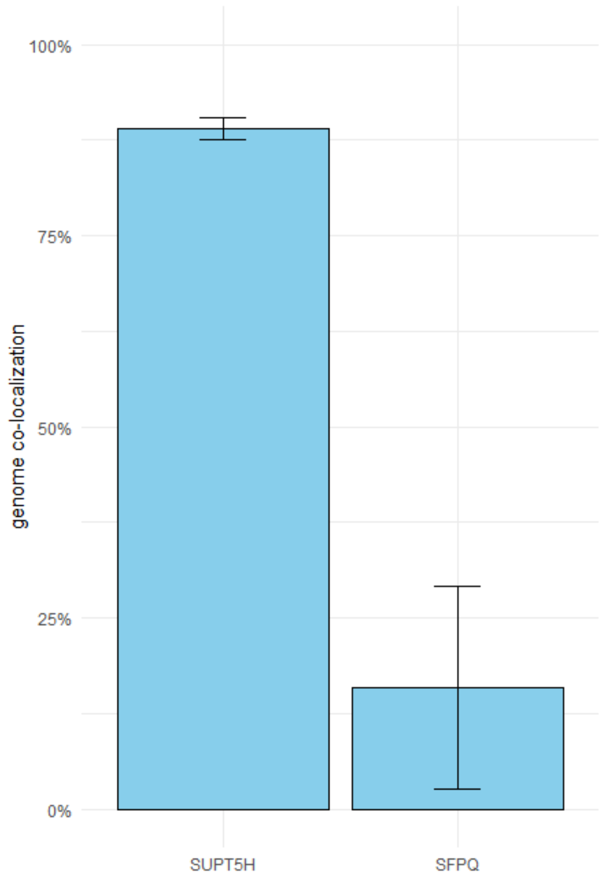


**Supplementary Figure 3: Quantification of genome co-localization.** Based on images from Figure 4, the proportion of viral DNA foci at locations of protein-specific signal was quantified. Protein-specific intensity thresholds were set by the median intensity at three randomly selected nuclear positions. Co-localization events were defined as signals exceeding the respective thresholds, and the proportion of colocalized spots relative to DNA signals was calculated. A total of 95 and 118 genomes were analyzed for SUPT5H and SFPQ, respectively.
